# Interpretation of ‘Omics dynamics in a single subject using local estimates of dispersion between two transcriptomes

**DOI:** 10.1101/405332

**Authors:** Qike Li, Samir Rachid Zaim, Dillon Aberasturi, Joanne Berghout, Haiquan Li, Francesca Vitali, Colleen Kenost, Helen Hao Zhang, Yves A. Lussier

**Author notes:** Contributed equally (Qike Li, Samir Rachid Zaim).

## Abstract

Calculating **<u>D</u>ifferentially <u>E</u>xpressed <u>G</u>enes (DEGs)** from RNA-sequencing requires replicates to estimate gene-wise variability, infeasible in clinics. By imposing restrictive transcriptome-wide assumptions limiting inferential opportunities of conventional methods (edgeR, NOISeq-sim, DESeq, DEGseq), comparing **two <u>c</u>onditions <u>w</u>ithout replicates** (**TCWR**) has been proposed, but not evaluated. Under TCWR conditions (e.g., unaffected tissue vs. tumor), differences of transformed expression of the proposed **individualized <u>DEG</u>** (**iDEG)** method follow a distribution calculated across a local partition of related transcripts at baseline expression; thereafter the probability of each DEG is estimated by empirical Bayes with local false discovery rate control using a two-group mixture model. In extensive simulation studies of TCWR methods, iDEG and NOISeq are more accurate at 5%<DEGs<20% (precision>90%, recall>75%, false_positive_rate<1%) and 30%<DEGs<40% (precision=recall∼90%), respectively.

The proposed **iDEG** method borrows localized distribution information from the same individual, a strategy that improves accuracy to compare transcriptomes in absence of replicates at low DEGs conditions. http://www.lussiergroup.org/publications/iDEG

## Introduction

Precision medicine aims to deliver “the right treatments, at the right time, to the right person”Kaiser [1]. However, clinical research, medicine, and pharmacology need new tools to achieve that goal. The prevailing system of one-size-fits-all drug development has led to the ten top-grossing USA drugs being ineffective for more than 75% of users^2^, and these patients typically cannot be identified until after therapeutic failure has occurred. The success of precision medicine hinges on identifying the precise aberrant mechanisms at play during an individual’s disease course^3^ to optimize treatment based on that individual’s biology.

Single-subject RNA sequencing (**RNA-Seq**) analysis considers one patient at a time, with the goal of revealing an individual’s altered transcriptomic mechanisms. Relative to traditional cohort-based analyses, a major challenge of single-subject RNA-Seq analysis is the estimation of gene expression variance which is required to identify differentially expressed genes (**DEGs**). In cohort-based methods, gene variance is calculated across a heterogenous set of samples, and the statistical methods employed leverage and rely on those replicates. However, they also emphasize consistent and average responses which may not accurately represent a single patient when the disease is heterogenous or stratified. Alternatively, the variance can be assessed between two conditions in one subject and three replicates. Yet, obtaining sufficient isogenic replicates for one subject to answer more precision questions poses a major difficulty due to (i) limited tissue availability, (ii) the risks associated with invasive tissue-sampling procedures, and (iii) general costs and inefficiencies with the current technology. Even though there is a great body of work for identifying DEGs in RNA-Seq data^4-8^ and frameworks for N-of-1 studies either for a single analyte or by pooling gene products in pathways^9-13^, to the best of our knowledge, no methods have been designed or validated at the gene level to determine the effect size and statistical significance of a single-subject, single RNA-Seq studies in **two conditions without replicates** (**TCWR**)^14^. Strategies to implement standard RNA-seq analysis methods for comparing TCWR have been proposed in the respective methods’ publications without comprehensive evaluation. Typically, these standard methods, usually requiring large cohorts, have been adapted to identify DEGs in TCWR by imposing restrictive transcriptome-wide distribution assumptions, thus limiting localized inferential opportunities.

Three critical obstacles hinder the analysis of single-subject TCWR studies. These include i) *patient-level inferential capability* in absence of biological replicates, ii) sensitivity to *fold-change inflation* in low-expression genes, and iii) *rigid parametric data assumptions* for variance estimation. To overcome the current technical limitations in analyzing RNA-Seq data, we propose a new method that borrows localized information across different genes from the same individual using a partitioned window to strategically bypass the requirement of replicates per condition: **iDEG** (**i**ndividualized **<u>D</u>**ifferentially **<u>E</u>**xpressed **<u>G</u>**enes). iDEG applies a localized variance-stabilizing transformation to estimate a gene’s distribution that borrows information from genes with similar baseline expression. While variance-stabilizing transformation has been previously used to identify DEGs across a large number of subjects or replicates, our approach differs from these since it has been developed to be applied directly on two paired transcriptomes from a single subject by computing the localized dispersion parameters in different windows of genes with similar expression at the baseline.

In this work, we evaluated the performances of iDEG and other four standard approaches applied to single-subject TCWR studies (edgeR, NOISeq-sim, DESeq, DEGseq). We also designed simulation studies under several conditions to stratify the range of applicability of our proposed strategy, which could eventually complement other RNA-seq analyses in TCWR studies. This study demonstrates the utility of variance-stabilizing transformations within subject in absence of replicates in two conditions, which is distinct from previous implementations of variance-stabilizing methods conducted across replicates or subjects.

## Methods

### The iDEG algorithm: iDEG

The iDEG algorithm (**Figure 1**) is an easy-to-implement, single-command function written in R^15^ with a computation speed of one second for identifying a subject’s DEGs on 8GB Ram computer. The subsequent sections expand on the main iDEG steps shown in **Figure 1**.

**Figure 1.**
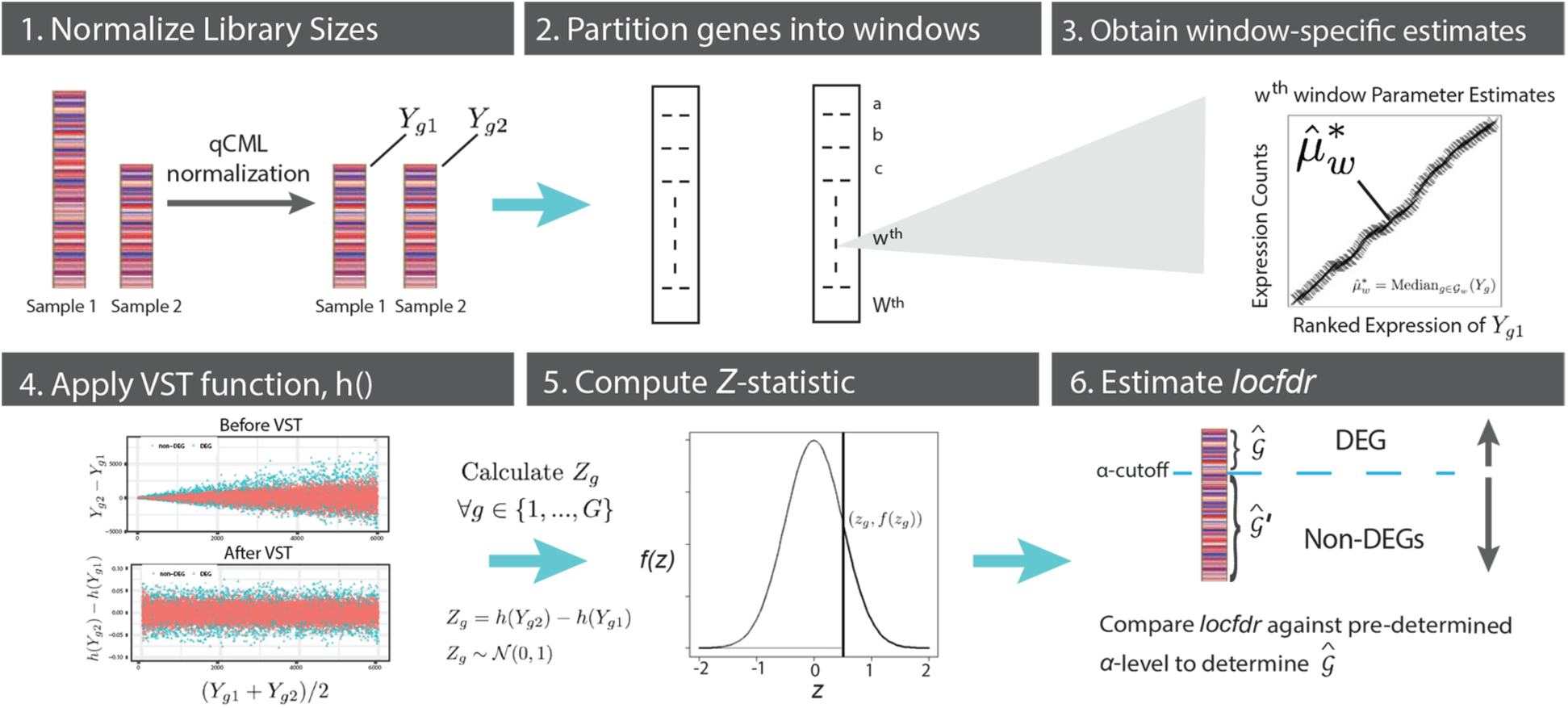
The iDEG Algorithm. 1) Normalize unequal library sizes if necessary. 2) Partition transcriptome into percentile-based windows using ranked baseline expression. 3) For each window: estimate mean expression, variance, and dispersion parameters. 4) Apply the Variance Stabilizing Transformation (**VST**) for each gene expression count. 5) Calculate the standard normal summary statistic “*Z*_*g*_” for each gene expression count “*g”*. 6) Determine the identified DEG set 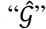 based on a pre-determined ***α-***cutoff.

### Modeling read counts via a re-parameterized Negative Binomial (NB) distribution

We model read counts *Y*_*gd*_ as following a re-parameterized negative binomial distribution with mean *μ*_*gd*_ and dispersion *δ*_*g*_. Thus, *Y*_*gd*_ ∼ *NB*(*μ*_*gd*_, *δ*_*g*_) with the following probability mass function, mean, and variance, respectively. Since for any subject, both transcriptomes are sequenced separately, they are treated as conditionally independent, conditional on the subject^*^.

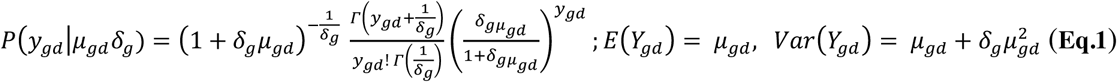

To identify DEGs from a pair of transcriptomes, we must test multiple hypotheses *H*_0_: μ_g1_ = μ_g2_, *g* = 1, …, G, where μ_*g*1_ and μ_*g*2_ are the theoretical mean expression levels for each gene “*g”* in sample 1 and sample 2, respectively. We define the DEG set by 𝒢 = {g: μ_*g*1_ ≠ μ_*g*2_, *g* = 1, …, G} and its set-theoretic complement of non-differentially expressed genes, or “null gene set” by 𝒢′ = {1, …, G}\ 𝒢. In presence of replicates, each hypothesis can be tested with a two-sample comparison, using Welch’s t-test statistic: 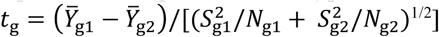 were 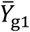 and 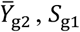 and *S*_g2_, and *N*_g1_ and *N*_g2_ are each groups’ respective sample mean, standard deviation, and size. However, when there is only one observation for *Y*_g1_ and one for *Y*_g2_, neither 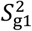 nor 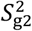 are computable. We thus propose iDEG: an algorithm that transforms *Y*_g1_ and *Y*_g2_, such that a simple function of the transformation allows for modeling all genes with the same distribution. This is done by pooling the genes together and estimating their common variance, hence bypassing the single-subject, single-replicate limitation.

### Normalize read counts with unequal library sizes (Figure 1, Panel 1)

In practice, unequal DNA library sizes may exist; thus, the first step is to normalize library sizes if necessary. We use the quantile-adjusted conditional maximum likelihood (**qCML**) procedure by Robinson and Smyth^16^, for normalization, and subsequently, iDEG is applied.

### Partition genes into windows to estimate local mean and variance (Figure 1, Panel 2)

Marioni et al. demonstrated the aptness of using expression means to estimate a gene’s variance^17^. Therefore, by extension, in iDEG we assume that genes of comparable expression levels are assumed to behave similarly (genes with similar means share similar variances). Thus, after normalization, the next step is to group genes into *W* non-overlapping windows of similar expression levels to approximate each window’s local mean and variance parameters. In the re-parameterized NB distribution, the variance of a given gene, g, is a function of its mean, *μ*g, and dispersion, *δ*g. Thus, the genes are partitioned to obtain their local, window-specific parameters. We define the *w*^*th*^ window by the (*w* - 1)^*th*^ and (*w*)^*th*^ percentiles, for *w=1*,…,*W*. 𝒢_*w*_ = {*g*: (*w* - 1)^*th*^ *percentile of Y*_g1_ < *Y*_g2_ < *w*^*th*^ *percentile of Y*_g1_}. To provide robust parameter estimates, we recommend a large positive integer, *W*, so that each window contains between 150 and 200 genes. However, the final predictions are not overtly sensitive to the choice of *W* (< 10% difference, data not shown – available upon request).

### Compute each window’s parameters (Figure 1, Panel 3)

As seen in (**Eq. 1**), 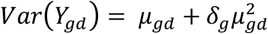. Therefore, when estimating variance locally, we are actually estimating the dispersion parameter *δ*_*g*_ for each gene count. This local estimation provides a more numerically fair evaluation of genes as it allows for comparisons relative to their mean expression counts. Particularly, it enables a better estimation of dispersion and variance parameters for genes with extremely high or low expression counts, since these genes are grouped together into windows, and share window-level parameter estimates in order to over-inflate or deflate their variability by averaging it out across the entire transcriptome. This is done in effort to mitigate challenges with making DEG calls in lowly and highly expressed genes. This value is required for the variance-stabilizing transformation (**VST**) h calculation (**Eq. 3**; **Figure 1 Panel 4**). However, variance cannot be estimated when only a single observation is available. For RNA-Seq data analysis, one common assumption is that the dispersion *δ*_g_ is equal across samples 1 and 2, and that dispersion is a function *q* of the mean, μ_*g*_^18-20^. Thus, in the absence of replicates, we partition genes into small windows to estimate the **functional mean-dispersion relationship**, *δ*_g_ = *q*(*μ*_g_), and hence the variance. We propose a two-step nonparametric procedure to obtain: (i) an initial estimate of *δ*g by pooling genes locally; and (ii) a refined estimate of *δ*_g_ by estimating *q(μ*_*g*_) with a smooth curve-fitting technique. In this approach, all non-differentially expressed (null) genes belonging to the same window 𝒢_w_ roughly have the same mean 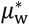 and the dispersion value 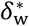. Thus, 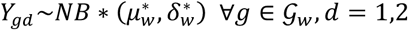 where 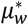 and 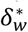 are the window-specific mean and dispersion values for null genes in 𝒢_*w*_, while *d* specifies if the count comes from sample 1 or 2.

The initial window estimates 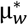 and 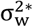 as 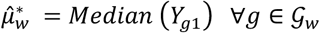 and 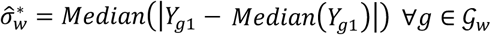, respectively. Since 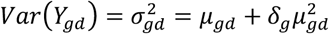, we estimate 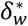 with 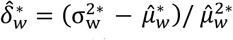, for all windows *w* = 1, …, *W*. To further improve the estimate of the dispersion parameter, *δ*_*g*_, a smoothing spline technique is used to fit a functional mean-dispersion relationship, *δ*_*g*_ = q(μ_*g*_), by solving the following optimization problem:

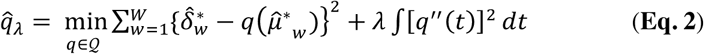

where 𝒬 is the second-order Sobolev space on [0, 1] containing *q*, and λ is a smoothing parameter (selected via generalized cross validation)^21^. After the fitted curve 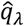 is obtained as in **Eq. 2**, the refined estimate of 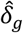 is computed as 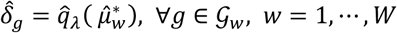. **Figure 2** illustrates the functional mean-dispersion relationship and calculation.

**Figure 2.**
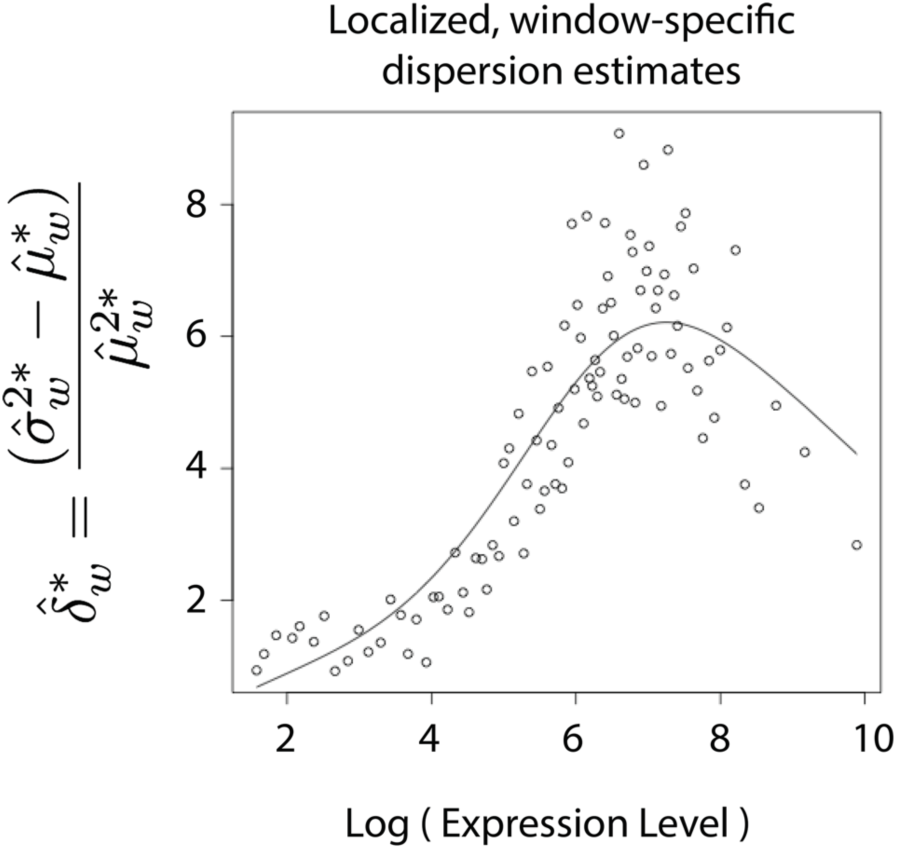
Localized, window-specific dispersion estimates as a function of log (mean expression). iDEG partitions the transcriptome into W equal-sized genomic windows of similar expression size and then calculates the over dispersion parameter relative to the gene’s mean expression. The number of windows W is a parameter in the iDEG function-call and should be empirically calculated relative to the transcriptome size. After conducting a few numerical studies, we recommend setting W=100 in order to allow for over dispersion estimates of highly and lowly expressed genes to be representative of their groups.

Summarizing, we get:

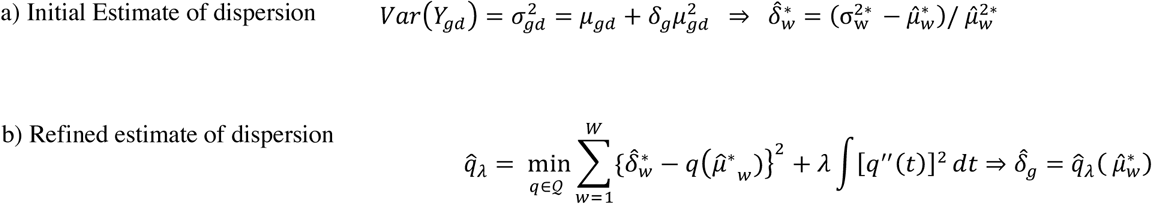

These equations come from the fact that in the negative binomial, variance (equation a) is a function of both mean and dispersion. So, the above equation (a) is rewritten version of the typical variance equation, with the stars and hat superscripts denote that it is now an estimate of the theoretical values for each partitioned window, w. As shown in (b), once window-level parameters are estimated (e.g., 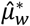), then a window-level dispersion parameter is estimated for all genes in that window (e.g., 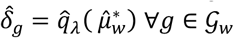), by fitting refined, functional estimate of dispersion.

### Apply the Variance Stabilizing Transformation (h(y_gd_)) to each gene (Figure 1, Panel 4)

After fitting *δ*_*g*_, we apply the variance stabilizing transformation *h* to the counts, *Y*_*gd*_ ∀*g* ∈ 𝒢_*w*_:

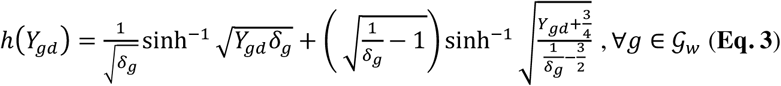

This transformation^22^ of the expression *Y*_*gd*_ in each window *w*, results in an approximately constant variance across all windows of the transcriptome (**Figure 3**), regardless of the expression mean, μ_*gd*_. That is 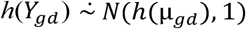, where *d* specifies if the count comes from sample 1 or 2. Therefore, the difference of the two independent normal random variables (e.g., *h*(*Y*_*g*1_) - *h*(*Y*_*g*2_)) approximately follows a common normal distribution with mean 0 and a constant variance: 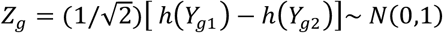. We suggest replacing 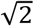 by a robust estimate of standard deviation (e.g., median absolute deviation)^23^. In most single-subject analyses, the estimated dispersion parameter, 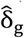, is small, but when 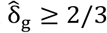, the VST *h*(*Y*_*gd*_) is not numerically stable. To avoid this issue, we suggest replacing *h* with *h*^* 24^, where 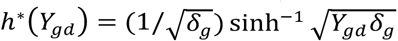, *g* = 1,…,*G*; *d* = 1,2. If a negative value of 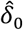 is obtained, we conservatively set it to zero to assume a larger variance.

**Figure 3.**
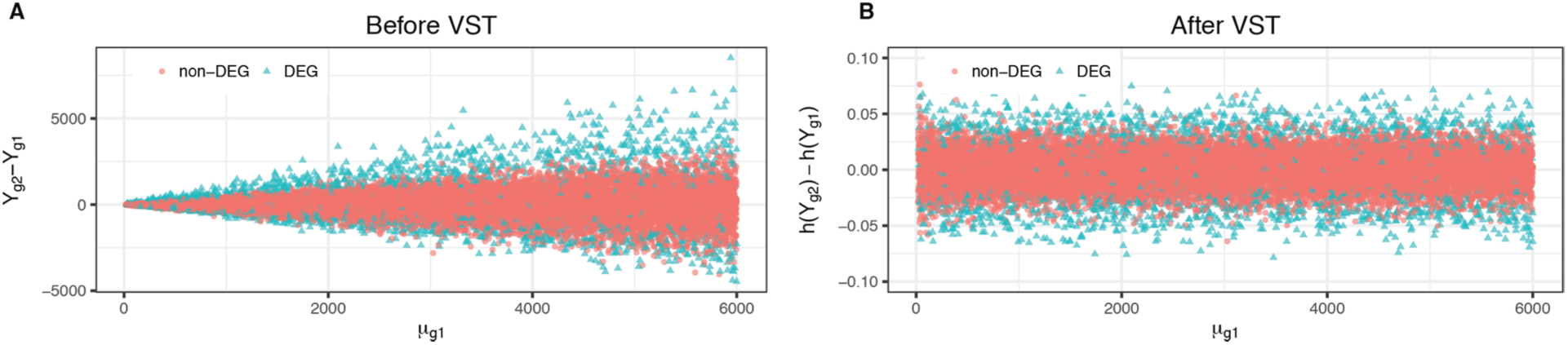
Variance Stabilizing Transformation (VST). **Panel A** depicts the raw difference *D*_*g*_ = *Y*_*g*1_ − *Y*_*g*2_ for 20,000 simulated genes (Methods Simulations), suggesting that the variance of D_g_ increases as the mean μ_g1_ increases; hence, there is no uniform cutoff to differentiate DEGs and null genes. **Panel B** illustrates that, for null genes, VST makes the variance of 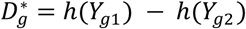 constant regardless of mean *μ*_*g*1_.

### Compute the summary statistic for each gene (Figure 1, Panel 5)

In the context of noisy data and large-scale inference, performing individual tests neglects the parallel structure of RNA-Seq data. Moreover, actual data mean and variance may not be close to their theoretical values of 0 and 1 due to various reasons (e.g., correlation across genes, correlation between samples, or failed mathematical assumptions)^25^. Therefore, we estimate an empirical null distribution *N*(μ_0_, *σ*_0_) to test these individual hypotheses.

Since differentially and non-differentially expressed genes generally follow different distributions, the probability density function of *Z*_*g*_, *f*(*z*), is naturally modeled by a two-group mixture: *f*(*z*) = *π*_0_*f*_0_ (*z*) + *π*_1_*f*_1_(*z*). Here, *f*_0_ and *f*_1_ are the probability density functions of genes in 𝒢′ and in 𝒢, while *π*_0_ and *π*_1_ = 1 – *π*_0_ are their respective membership proportions. We assume a normal distribution following previous work from Dean and Raftery^26^ that applied a two-group mixture model to identify differentially expressed genes, assuming a normal distribution for the null genes and a uniform distribution for the DEGs. However, we relax their assumptions for the marginal distribution and assume an exponential family. We approximate *f* (*z*) using a smooth K-parameter exponential family distribution, 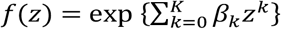, and estimate the parameters (*β*_0_, *β*_1_, · · *·, β*_*K*_)^*T*^ using Efron’s approach^27^

### Estimate the local false discovery (locfdr) for each and identify DEGs (Figure 1, Panel 6)

Finally, to control the false discovery rate (**fdr**), we adopt Efron’s idea^28-30^ to estimate the local fdr (***locfdr***) using the R package *locfdr* and estimate *π*_0_, *f*_0_(*z*) by maximum likelihood. Efron et al.^31^ have shown *locfdr*’s close connection to the BH false discovery rate procedure^32^; therefore, after estimating *locfdr(zg)*, it identifies differentially expressed genes by comparing *locfdr(z*_*g*_) to a pre-specified **α**-cutoff value. The final set of differentially expressed genes identified by applying the iDEG procedure is denoted by 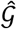.

### Simulations (Figure 4)

To compare the performance of iDEG to existing methods – including edgeR^16^, DEGSeq^8^, DESeq^19^, and NOISeq^6^ – extensive numerical studies were conducted assuming that RNA-Seq data follow the NB distribution with a varying dispersion parameter *δ*_*g*_. Of note, these methods assume the NB distribution for data, which is used in the simulation; except for NOISeq that is nonparametric and DEGseq which assumes a binomial distribution. Baseline (normal tissue) and case (tumor sample) transcriptomes are both simulated and assumed to contain G = 20,000 genes; the library size of one transcriptome is 1.5 times larger than the other one. The single-subject RNA-Seq datasets are simulated with different percentages of DEGs, including DEG percentage = 5%, 10%, 15%, 20%, 25%, 30%, 35%, and 40%, and also with different window sizes W =10, 100, and 1000 (data not shown for window-level experiments). Each experiment is repeated 1000 times, across each of these simulation conditions. *Y*_*g*1_ ∼ NB(μ_*g*1_, *δ*_*g*_) and *Y*_*g*2_ ∼ NB(μ_*g*2_, *δ*_*g*_), where *μ*_*g*1_follow a discrete uniform over the range *B* = {5, 6, …, 10,000}, and the dispersion parameter *δ*_*g*_ has been set to *δ*_*g*_ = 0.005 + 9/(μ_*g*1_ + 100), per Anders and Huber^19^. Probabilities for gene expression means, μ_*g*1_, are sampled from: 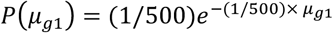, *g* = 1, …, 20000. For the case transcriptome, we set μ_*g*2_*=* μ_*g*1_ for *g* ∈ 𝒢 ′ and *μ*_*g*2_ = *d*^*s*^ *μ*_*g*1_ for *g* ∈ 𝒢, where *s* = (−1)^*b*^ and b ∼ Bernoulli(0.5) is a random variable, and 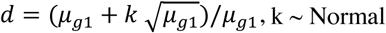 (4,1). Here, s indicates increasing expression (s = 1) or decreasing expression (s = −1) of a gene in the case transcriptome relative to baseline. Finally, for each gene g, we simulate one observation for *Y*_*g*1_ and *Y*_*g*2_ respectively and test the hypothesis μ_*g*1_ = μ_*g*2_. At each iteration, a baseline and a case transcriptome are generated to simulate a distinct RNA-Seq dataset. Methods are assessed by their Precision, Recall, and FPR, and *F*_1_ score, *F*_1_ = (2 · (precision * recall)/(preci*s*ion + recall). The average number of identified DEGs is also reported. Of note, we excluded from the comparison GFOLD, a standard approach that can be applied to TCWR studies, as it only ranks genes without providing a measure of significance, thus prohibiting the accurate comparison with the remaining techniques using precision-recall curves or ROC curves.

**Figure 4.**
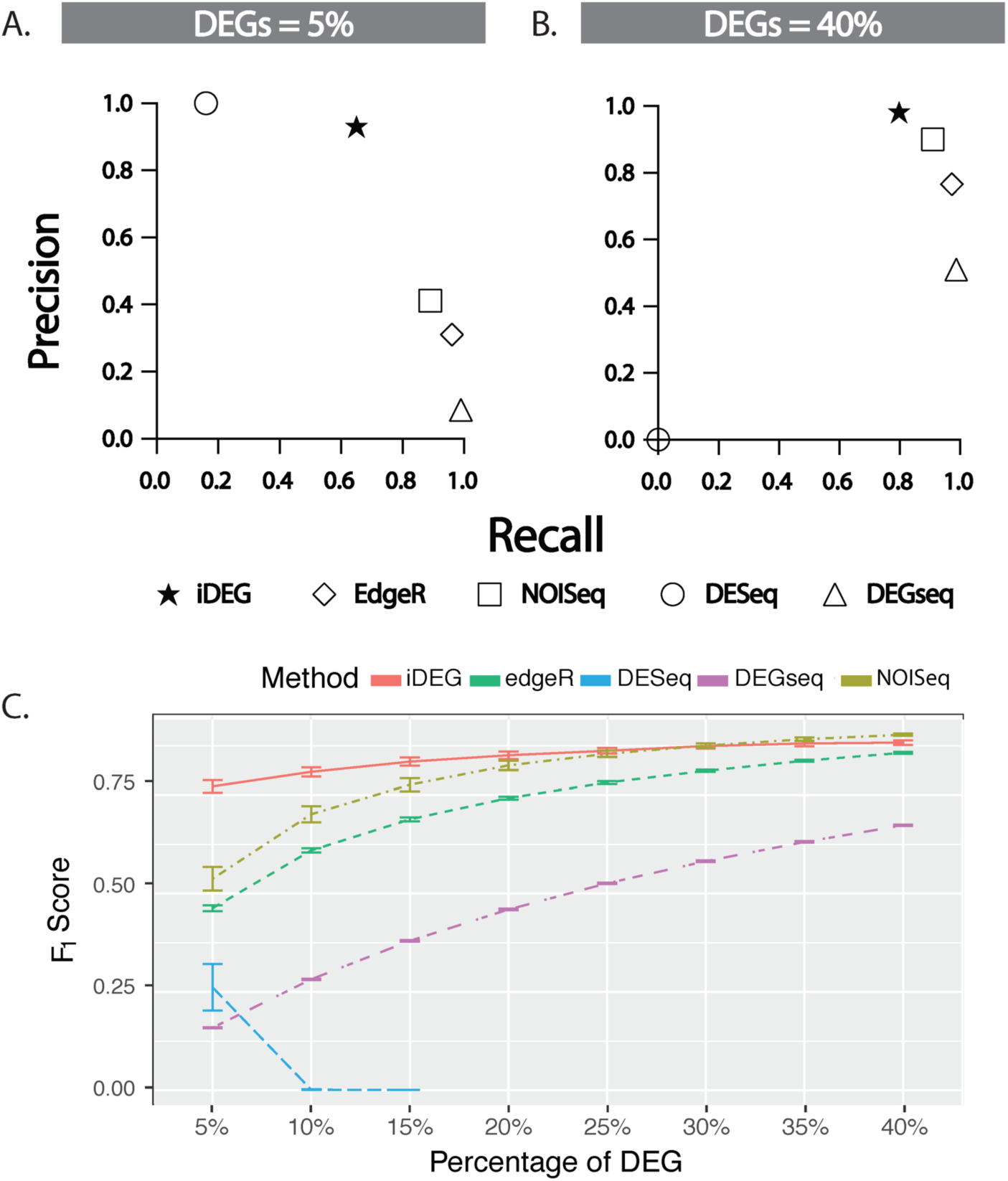
Performance results. NOISeq-sim’s and iDEG’s F_1_-scores are more accurate than that of other methods at 5%<DEGs<20% (iDEG) and 30%<DEGs<40% (NOISeq-sim). At DEG=5% and FDR<10%, iDEG provides an interesting compromise between precision and recall, while NOISeq provides a better compromise when the percentage of DEGs is higher than 30%. **Panels A & B**. Precision recall curve at 10% FDR for 1,000 and 8,000 seeded DEGs among 20,000 transcripts, respectively. **Panel C.** F_1_ scores. Average F_1_ scores resulting from 1,000 repeated experiments with vertical bars representing one standard deviation.

## Results

**Figure 4** depicts the accuracies obtained in the simulations while **Table 1** contextualizes each method’s performance relative to the number of DEG calls and the number (and %) of genes seeded as DEGs. Of note, we have also conducted complementary analyses with a Poisson distribution and showed similar ordering of accuracies between the evaluated methods (data not shown). For windows size of 10, 100, and 1000 the accuracies of the simulations remained consistent (data not shown) and opted for setting W = 100 to balance computation time and parameter estimation robustness. As seen in Table 1, NOISeq-sim, edgeR, DEGseq attain a high precision (defined as > 90% precision) across all simulation conditions (5% through 40% DEGs seeded) at the expense of lower recall and a large number of false positives. For example, as seen in **Table 1**, 5% DEGs, NOISeq-sim, edgeR, and DEGseq all result in a larger of false positives than there are actual seeded genes. Conversely, iDEG attains a high precision (defined as >90%) at the expense of making a smaller number of DEG calls, thus attaining lower recall. The F1 score shown in Fig 4-C is the harmonic mean of the Precision and Recall metrics, aggregating the precision-recall trade-offs made by individual techniques into a single technique. Although iDEG never attains as high a recall as DEGseq, edgeR, and NOISeq-sim, it better balances its precision-recall trade-off into a higher overall F1 score at FDRs<20%, while NOISeq does better at FDR>30%, and the two methods show similar F1 scores at 20%<FDR<30%. Of note, DESeq failed to make any DEG calls across the majority of the simulation conditions (since it either produced “0” or “1” fdr-adjusted probability predictions), preventing us from evaluating their performance at any reasonable false discovery cutoff.

**Table 1.** Performance results of TCWR simulations. At different percentage of DEGs in TCWR simulations, distinct methods obtain the best precision and recall, with iDEG, NOISeq and edgeR producing the best combinations of precision and recall. Of note, edgeR, DEGseq, and DESeq were not designed nor validated for studies without replicates; however, their authors proposed to utilize them in these conditions by defining specific parameters. NOISeq-sim offers high recall and precision with DEGs=40% i.e. when 8,000 genes are dysregulated among 20,000. On the other hand, iDEG obtains high precision with moderate to high recall in all conditions. EdgeR provides moderate precision with very high recall for DEGs>20%.

| Proportions of DEGs seeded | Method | Precision | Recall (TPR) | FP | Predicted DEGs |
| --- | --- | --- | --- | --- | --- |
| **1,000** out of 20,000<br>(5% of genes) | **iDEG** | **0.93** | 0.65 | 57 | 700 |
|  | edgeR | 0.31 | 0.96 | 2,090 | 3,119 |
|  | NOISeq-sim | 0.412 | 0.89 | 1140 | 2,163 |
|  | DESeq | 1.0 | 0.16 | 0 | 162 |
|  | **DEGseq** | 0.086 | **0.99** | 10,450 | 11,397 |
| **4,000** out of 20,000<br>(20% of genes) | **iDEG** | **0.97** | 0.76 | 112 | 3,136 |
|  | edgeR | 0.60 | 0.97 | 2,560 | 6,405 |
|  | NOISeq-sim | 0.747 | 0.91 | 1120 | 4,897 |
|  | DESeq | Not applicable | 0 | 0 | 0 |
|  | **DEGseq** | 0.3 | **0.99** | 9,280 | 13,160 |
| **8,000** out of 20,000<br>(40% of genes) | **iDEG** | **0.98** | 0.80 | 120 | 6,484 |
|  | edgeR | 0.77 | 0.97 | 2,400 | 10,145 |
|  | NOISeq-sim | 0.893 | 0.91 | 840 | 8,177 |
|  | DESeq | Not applicable | 0 | 0 | 0 |
|  | **DEGseq** | 0.51 | **0.99** | 7,560 | 15,455 |

## Discussion

No single method has emerged as the optimal approach for all conditions. Low expression levels are extremely susceptible to unstable fold-change estimation, as a 5-fold increase from 2 to 10 counts on a dynamic range of 0 to 100,000 should not be treated equivalently to that between 10,000 and 50,000. Standard practice filters out genes with counts below a certain threshold (typically 5 or 10). However, this solution does not address fold change (FC) inflation above the threshold (e.g., FC>2 at 15 counts), nor how to compare distinct FCs at different expression levels. Alternatively, favoring absolute count difference to identify DEGs leads to a systemic bias towards genes with high expression. Conversely, favoring FC results in a systemic bias towards lowly expressed genes. Either of these solutions yields higher false positive rates. For DEGs<30%, the variance stabilization within partitioned windows proposed in iDEG is shown to address this dilemma of dealing with fold change inflation by comparing FC values relative to their expression levels, perhaps because conventional approaches impose stringent data assumptions that may compromise downstream inferential processes.

As we proceeded to validating iDEG in biologic or clinical datasets, a review of literature identified few candidate datasets that comprised targeted mutations over an isogenic background and yield high DEG rates (e.g., DEGs>50%) that did not reflect rates expected in clinical care. In addition, the state of the art in generating reference standard consisted in comparing one method against itself as the overlap of DEGs across conventional methods was low in spite of 30 replicates in isogenic conditions. Because of these two considerations, we decided to publish the results of a comprehensive improvement in reference standard generation as a companion paper ^33^. We have thus generated multiple distinct reference standards (one per conventional method) and developed a “fair” evaluation of methods to identify DEGs in paired conditions without replicates using *biological datasets* (each method is compared to all other methods but not itself)^33^. This companion biological paper^33^ is limited to datasets with high DEGs as no reference datasets were available for low DEGs conditions, while the current simulation explores both low and moderate DEGs levels.

We note several limitations to the current study. First, conventional techniques were not explicitly designed for absence of replicates and are tested in those conditions. In addition, each method assumes some distribution (DEGseq assumes a binomial distribution; iDEG, edgeR, and DESeq assume a Negative Binomial distribution; and NOISeq is non-parametric). Since the distributional form in real-data is never truly known (only approximated), simulating a transcriptome necessarily entails distributional assumption in every simulation study, which limit its generalizability to real studies and inherently may favor some methods over the others. Furthermore, not all DEG detection techniques can actually produce a DEG call in TCWR. This limits the number of techniques available for comparison in our simulation study. Moreover, as seen with DESeq, DEG techniques designed for replicated studies are not necessarily fully operational or effective in TCWR, therefore it is not necessarily recommended to pick an arbitrary DEG technique and use it in non-replicated TCWR studies. We conducted the simulations against these methods to illustrate the need for new approaches to study single-subject transcripts in TCWR conditions. In addition, a true gold standard to evaluate iDEG and other methods is not as simple as obtaining replicates and running conventional methods as pointed out by recent papers^34, 35^.

## Conclusion

Over the past decade, state-of-the-art techniques in RNA-Seq data analysis have delivered powerful new tools^14^ for extending large-scale inference to small-sample settings. The primary goal of iDEG is not to replace these, but rather to expand the scope of RNA-Seq studies into the single-subject, single-replicate realm and provide novel research opportunities and test methods for controlling fold change inflation at low expression ranges. In iDEG, we have shown the novelty of window partitioning to borrow localized distribution information across genes, and its improved accuracy over alternate methods in low DEG conditions (DEG<20%). Furthermore, this approach could potentially be applied to improve the accuracy of existing parametric and non-parametric differential expression tools. In future studies, we envision to i) extend the window partitioning component of iDEG into other techniques, ii) to locally identify differentially expressed pathways (by incorporating ontologies and knowledge graphs), and iv) to apply it to other ’omics measures, (e.g., metabolomics, proteomics, etc.).

## List of Abbreviations

DEGs: differentially expressed genes
FC: fold change
FCI: fold change inflation
FDR: false discovery rate
iDEG: individualized Differentially Expressed Genes
*locfdr*: local false discovery rate
NB: Negative Binomial distribution
RNA-Seq: RNA Sequencing
TCWR: two conditions without replicates
VST: Variance Stabilizing Transformation

## Declarations

### Availability of data and material

Software is available at http://www.lussiergroup.org/publications/iDEG

### Authors’ contributions

Conceived the study: QL, HHZ, YAL. Experimental design and analysis: SRZ, QL, DA, YAL, HHZ. Manuscript writing: SRZ, QL, YAL, JB Figures: SRZ, CK, YAL. Interpretation: SRZ, QL, DA, JB, FV, HL, YAL. All of the authors have read and approved the final manuscript.

## Footnotes

* A note on notation, since iDEG models each patient’s paired transcriptome individually, the subscripts for each subject are omitted since only a subject is handled at a time in any given calculation.

